# Evaluation of a novel Tc-24 recombinant antigen ELISA for serologic testing for *Trypanosoma cruzi* in dogs

**DOI:** 10.1101/2024.02.12.579969

**Authors:** Rojelio Mejia, Guilherme G. Verocai, Ilana A. Mosley, Bin Zhan, Lindsey Vongthavaravat, Rachel E. Busselman, Sarah A. Hamer

**Affiliations:** Department of Pediatrics - Tropical Medicine, Baylor College of Medicine, Houston, Texas; Department of Veterinary Pathobiology, College of Veterinary Medicine and Biomedical Sciences, Texas A&M University, College Station, Texas; Department of Veterinary Integrative Biosciences, College of Veterinary Medicine and Biomedical Sciences, Texas A&M University, College Station, Texas

**Author notes:** Corresponding author. 1102 Bates Ave, 0550.10, Houston, Texas 77030.

**Keywords:** Dogs, Chagas disease, Trypanosoma cruzi, Tc-24, Texas

## Abstract

Chagas disease is a parasitic infection caused by *Trypanosoma cruzi*. Diagnosis of chroni Chagas disease in dogs relies on limited serological test options. This study used a new Tc-24 recombinant antigen ELISA on an archival set of 70 dog serum samples from multi-dog kennel environments in Texas subjected to three existing Chagas serological tests. Tc-24 ELISA produced a quantitative result and could detect anti-*T. cruzi* antibodies in dogs with high sensitivity and specificity. Comparing individual tests to Tc-24 ELISA resulted in strong associations and correlations, which suggest that Tc-24 ELISA is a reliable and accurate diagnostic tool for dogs with a single test.

---

Chagas disease, or American trypanosomiasis, is an infectious tropical disease caused by the protozoan parasite *Trypanosoma cruzi*(1). *T. cruzi* infections in humans and other mammal hosts, including dogs, can occur via congenital transmission, blood transfusions/organ transplants, contact with infected triatomine feces, or ingesting an infected triatomine insect vector(1). It is ndemic to 21 countries across South and Central America, Mexico, and the southern United States(2). In humans, the diagnosis of chronic infection relies solely on serological tests. The diagnostic gold standard for diagnosing human patients with suspected chronic *T. cruzi* infection is the combining of two positive serological tests (enzyme-linked immunosorbent assay (3), hemagglutination inhibition assay [HAI], or indirect immunofluorescence [IIF]) and potentially a third test if the results are conflicting(4). However, existing serological tests for Chagas disease are prone to inconclusive results and exhibit poor concordance between different assay formats. A recent study showed that the performance of three commercial *T. cruzi* serological tests varied significantly when detecting different geographic *T. cruzi* strains, including 12% missed cases from Argentina, 21% missed cases from Honduras, and 72% missed cases from Mexico(5). These differences exist in both human and canine testing. Dogs are a known reservoir for *T. cruzi* across various endemic areas, particularly the southern U.S., including Texas, where infection prevalence between 20.3 – 31.6% and as high as 57.6% have been documented in domestic dogs and kennel environments, respectively(6, 7). Currently, there are limited immunodiagnostic tests specific for veterinary use. Therefore, many epidemiological studies in domestic and wild animal reservoirs rely on an IFA test, often used with commercial immunodiagnostic tests approved and intended for human use(8).

A concern surrounding Chagas diagnostics is that different diagnostic tests may perform differently in humans or animals infected with the seven known discrete typing units(9) and can potentially cross-react with related protozoans that may be co-endemic/or co-occur (e.g., *Leishmania*)(10). Our study aims to compare the performance of three commercial diagnostic tests to our Tc-24 recombinant antigen ELISA in samples of dogs from an endemic region. Tc-24 is a 24 kDa calcium-binding protein ubiquitously expressed in all stages of *T. cruzi*(11). It is an immunodominant protein during infection and conservative across all geographical strains of *T. cruzi* (97% homologous) but not present in other protozoa or helminths(12), suggesting that it can detect antibodies induced by all strains of *T. cruzi*. Our study examined 70 archival serum samples collected from dogs in ten large kennels in south Texas. The dogs were sampled up to three times each between May 2018 and September 2019 to initially collect data for a study to quantify the incidence of canine *T. cruzi* infections(6). Dogs enrolled in the study were between the ages of 13 months and 12.2 years (mean = 6.1 years; median = 6.1 years) and included hound dogs, Belgian Malinois, German Shorthair Pointers, Labrador Retrievers, Brittany Spaniels, and English Pointers. Samples were kept frozen at -80°C until processing.

A Tc-24 recombinant antigen ELISA protocol previously used for human samples (unpublished) was adapted for canine samples. Briefly, a capture ELISA was developed based on a *Strongyloides* antigen but using Tc-24 antigen at the same concentration instead of the *Strongyloides* antigen(13). A different secondary antibody was used to detect canine IgG, Rabbit anti-canine, Fc fragment, alkaline phosphatase at 1:2500 dilution (Jackson ImmunoResearch Laboratories, Inc. West Grove, PA). For the Tc-24 ELISA, the threshold of positivity was estimated using positives and negatives for the respected commercial tests and calculating the receiver operating characteristic curve (ROC). All Tc-24 ELISA plates were run on the same day to minimize inter-plate variability.

As previously reported(6), serum samples were tested for *T. cruzi* antibodies using three serological tests: Chagas Stat-Pak (‘SP’; ChemBio, Medford, NY, USA), Chagas *Detect Plus* Rapid Test (‘IB’; InBios International, Inc., Seattle, WA, USA), and an indirect fluorescent antibody (IFA) test. The Chagas Stat-Pak and Chagas *Detect Plus* are rapid tests that, while not labeled for use in dogs, are commonly used for research purposes to detect *T. cruzi* antibodies(8, 14, 15). For the current study, the strength of the response on the rapid tests was scored on a scale from 0-4, in which 0 was a negative result (no color development or when a very faint incomplete band developed) and scores of 1-4 were positive results with increasing intensity of the reaction line. The IFA was run at the Texas A&M Veterinary Medical Diagnostic Laboratory with endpoint titers determined. Both the IFA and Tc-24 ELISA produced a quantitative result. Results were compared using Mann-Whitney tests for nonparametric variables. Spearman correlation tests were used to determine relationships between continuous variables. Analysis of covariance was utilized for comparison of linear correlations. *P* < .05 was considered significant. These tests were performed using GraphPad Prism 8.2.1 (GraphPad Software, San Diego,California, USA).

Comparing individual tests to Tc-24 ELISA resulted in SP positives having higher means of Tc-24 OD than SP negatives (0.475 versus 0.098, p < 0.001) (Sensitivity 82.1%, Specificity 86.7%) (Figure 1A). IFA positives had higher means of Tc-24 OD than IFA negatives (0.607 versus 0.089, p < 0.001) (Sensitivity 87.5%, Specificity 91.2%) (Figure 1B). IB positives had higher means of Tc-24 OD than IB negatives (0.482 versus 0.091, p < 0.001) (Sensitivity 80.0%, Specificity 93.1%) (Figure 1C). SP results had an association with Tc-24 (p < 0.001) and a correlation of Spearman r = 1.0 (p = 0.0167), with higher SP values associated with higher Tc-24 results (Figure 2A). There was an association of IFA to Tc-24 (p < 0.001) and a correlation of Spearman r = 0.6273 (p = 0.044) (Figure 2B). The IB strips compared to Tc-24 also showed an association with Tc-24 (p < 0.001) and a correlation of Spearman r = 0.9 (p = 0.0833) (Figure 2C). Comparing any two positive commercial tests to the Tc-24 ELISA showed an association between positive versus negative results (p < 0.0001) with an 82.5% sensitivity and 89.7% specificity (results not shown). Comparing the highest IFA titers (>1280) of this study to Tc-24 (OD) resulted in (1.0 versus 0.089, p < 0.001) and 100% sensitivity and 94.1% specificity (Figure 3A). Similarly, comparing high (SP, IB) immunochromatographic values (4) to Tc-24 (OD) revealed (1.17 versus 0.039, p < 0.001) and 100% sensitivity and 95.8% specificity (Figure 3B).

**Figure 1.**
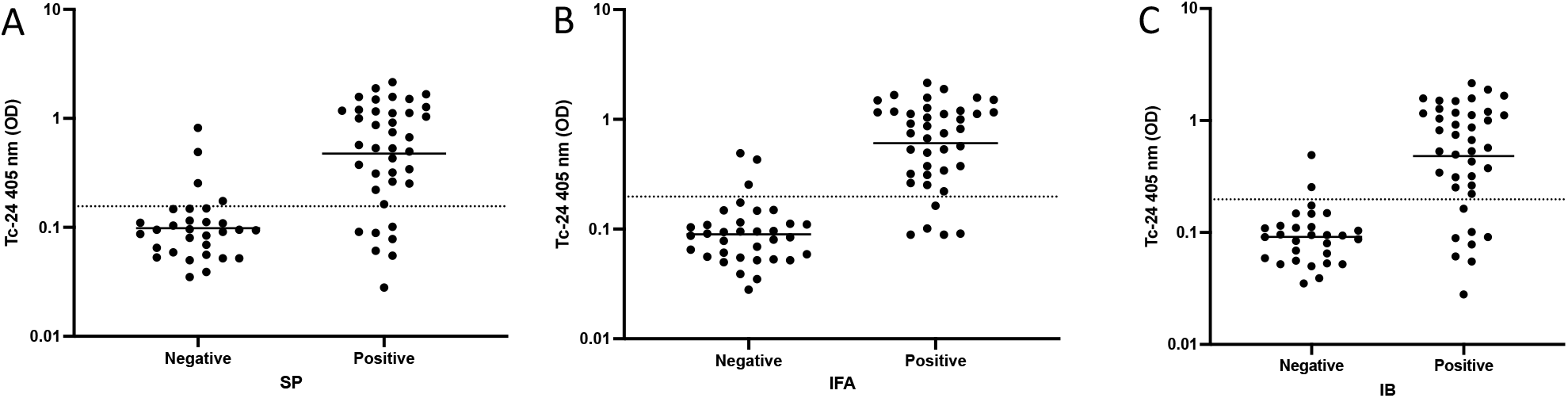
A) SP positives had higher Tc-24 OD than SP negatives (0.475 versus 0.098, p < 0.001). The threshold of positivity determined by the ROC curve was 0.156 OD (Sensitivity 82.1%, Specificity 86.7%). B) IFA positives had higher Tc-24 OD than IFA negatives (0.607 versus 0.089, p < 0.001). The threshold of positivity determined by the ROC curve was 0.198 OD (Sensitivity 87.5%, Specificity 91.2%). C) IB positives had higher Tc-24 OD than IB negatives (0.482 versus 0.091, p < 0.001). The threshold of positivity determined by the ROC curve was 0.198 OD (Sensitivity 80.0%, Specificity 93.1%).

**Figure 2.**
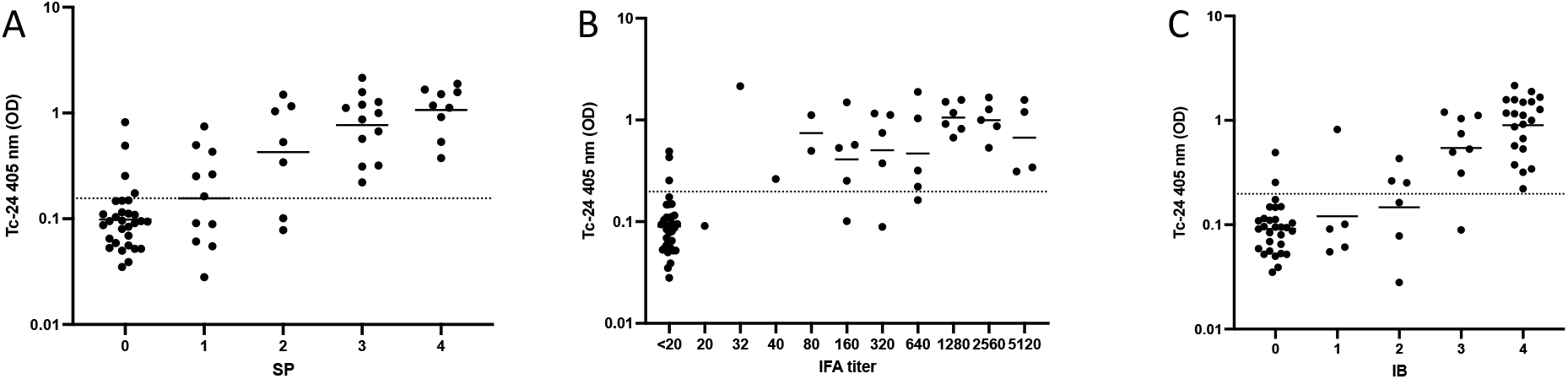
A) Tc-24 correlates to SP commercial testing (Spearman r = 1.00, p = 0.0167), and all groups were significantly different (ANOVA, p < 0.001). B) Tc-24 correlates to IFA commercial testing (Spearman r = 0.6273, p = 0.0440), and all groups were significantly different (ANOVA, p < 0.0001). C) Tc-24 correlates to IB commercial testing (Spearman r = 0.9, p = 0.0833), and all groups were significantly different (ANOVA, p < 0.001).

**Figure 3.**
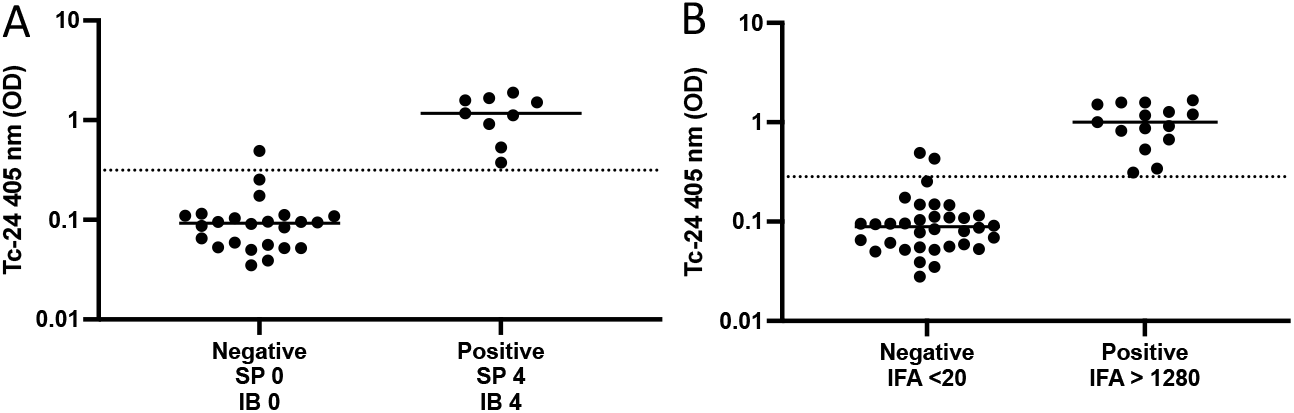
A) Comparing high (SP, IB) immunochromatographic values (4) to Tc-24 (OD) revealed (1.17 versus 0.039, p < 0.001) and 100% sensitivity and 95.8% specificity. B) Comparing the highest IFA titers (>1280) of this study to Tc-24 (OD) resulted in (1.0 versus 0.089, p < 0.001) and 100% sensitivity and 94.1% specificity.

Diagnosing Chagas disease in dogs is challenging due to *T. cruzi* biology and fluctuations in parasitemia throughout an infection(16). Efforts to improve veterinary Chagas diagnostics are underway; for example, the same archival canine serum was used to validate a multiplex microsphere immunoassay with promising results from a single assay(17), but this approach requires the use of the Luminex platform, which may not be readily available in veterinary diagnostic labs.

The results showed that SP and IB strips had a significant association and high correlation to Tc-24 ELISA. IFA also showed a significant association and moderate correlation to Tc-24 ELISA. Moreover, comparing any two positive commercial tests to the Tc-24 ELISA revealed a higher sensitivity and specificity than a single commercial test. Comparing high IFA titers and high immunochromatographic values to positive Tc-24 showed high sensitivity and specificity, respectively. These results indicate that Tc-24 ELISA may have the potential to be a new serological assay to diagnose *T. cruzi* infection in dogs with a single test.

The use of Tc-24 in diagnosing Chagas disease in dogs has not been widely studied. This study is significant as it suggests that Tc-24 recombinant antigen ELISA can potentially improve Chagas disease diagnosis in dogs. While this study provides promising results, several limitations need to be addressed. Firstly, the study had a small sample size, and the results must be replicated in larger studies. Secondly, the study only evaluated dogs from multi-dog kennel environments in Texas, and it is unknown whether the results can be generalized to other geographical locations or different populations of dogs. The study did not compare the Tc-24 recombinant antigen ELISA with other diagnostic assays for Chagas disease, such as Western blot or PCR-based tests. Lastly, the subjective grading of immunochromatographic tests may have introduced potential bias in the results.

## Summary

In conclusion, Chagas disease diagnosis in dogs is challenging due to the limitations of existing serological tests. This study suggests that Tc-24 recombinant antigen ELISA has potential as a new serological assay to aid in diagnosing *T. cruzi* infection in dogs with a singular test. These findings have implications as early diagnosis and treatment are crucial to preventing the development of chronic disease, reducing transmission, and broadening epidemiological surveillance in endemic areas.

## Acknowledgments

American Kennel Club Canine Health Foundation Grant No 02448 provided funding for dog sample collection. The funders had no role in study design, data collection and analysis, publication decisions, or manuscript preparation.

## Funding

NIH R21 1R21AI171477-01A1

## Author’s contributions

RM, GGV, IAM performed the Tc-24 ELISA. RM, BZ developed the Tc-24 ELISA. RM, GGV, SAH performed data analysis. All authors contributed to writing and editing the manuscript.

## Notes

### Competing Interest Statement

The authors have declared no competing interest.

